# Quadruplex qPCR for qualitative and quantitative analysis of the HIV-1 latent reservoir

**DOI:** 10.1101/641951

**Authors:** Christian Gaebler, Julio C. C. Lorenzi, Thiago Y. Oliveira, Lilian Nogueira, Victor Ramos, Ching-Lan Lu, Joy A. Pai, Pilar Mendoza, Mila Jankovic, Marina Caskey, Michel C. Nussenzweig

## Abstract

HIV-1 infection requires life-long therapy with anti-retroviral drugs due to the existence of a latent reservoir of transcriptionally inactive integrated proviruses. The goal of HIV-1 cure research is to eliminate or functionally silence this reservoir. To this end there are numerous ongoing studies to evaluate immunologic approaches including monoclonal antibody therapies. Evaluating the results of these studies requires sensitive and specific measures of the reservoir. Here we describe a relatively high throughput combined quantitative polymerase chain reaction (qPCR) and next generation sequencing method. Four different qPCR probes covering the packaging signal (*PS*), group-specific antigen (*gag*), polymerase (*pol*), and envelope (*env*) are combined in a single multiplex reaction to detect the HIV-1 genome in limiting dilution samples followed by sequence verification of individual reactions that are positive for combinations of any 2 of the 4 probes (Q4PCR). This sensitive and specific approach allows for an unbiased characterization of the HIV-1 latent reservoir.

**Summary:** HIV-1 cure research seeks to decrease or eliminate the latent reservoir. The evaluation of such curative strategies requires accurate measures of the reservoir. Gaebler et al. describe a combined multicolor qPCR and next generation sequencing method that enables the sensitive and specific characterization of the HIV-1 latent reservoir.

## Introduction

Like other retroviruses, HIV-1 integrates into the host genome where it is transcribed to produce infectious virions (Craigie and Bushman, 2012). Productive infection typically leads to cell death, however, in a small number of CD4^+^ T cells the integrated virus is silenced and becomes latent. Combination antiretroviral therapy (cART) is highly effective in suppressing HIV-1 infection and preventing disease progression, however, cART does not eliminate the virus due to the existence of the latent reservoir (Chun et al., 1997; Finzi et al., 1997; Wong et al., 1997). Longitudinal studies performed on individuals on cART indicate that the latent reservoir has a half-life of 44 months (Chun et al., 1999; Siliciano et al., 2003). Thus, treatment interruption leads almost invariably to rapid viral rebound and therapy with cART is required for the lifetime of the infected individual (Margolis and Archin, 2017; Sengupta and Siliciano, 2018).

An important goal for HIV-1 research is to achieve a functional remission or cure by decreasing or eliminating the latent reservoir, and a number of clinical trials have been designed to test new approaches to this problem (Caskey et al., 2019; Cillo and Mellors, 2016; Gruell and Klein, 2018; Margolis et al., 2017). The evaluation of HIV-1 curative strategies requires sensitive, specific and precise assays to quantify and characterize the latent HIV-1 reservoir. Yet, to date most approaches show major discrepancies in infected cell frequencies (Eriksson et al., 2013; Sengupta and Siliciano, 2018). These inconsistencies constrain the accurate assessment of HIV-1 cure efforts and could obscure a meaningful intervention (Henrich et al., 2017; Sengupta and Siliciano, 2018).

Here we report a relatively high throughput method for enumerating and characterizing intact latent proviral DNA by a combination of multicolor quantitative PCR (Q4PCR) and next generation sequencing. We compare the new method to quantitative and qualitative viral outgrowth assays (Q^2^VOA) and near full-length (NFL) sequencing on paired peripheral blood samples obtained at 2 time points from the same 6 individuals enrolled in a clinical trial that involved analytical treatment interruption (ATI) after infusion of a combination of two broadly neutralizing monoclonal antibodies (Lu et al., 2018; Mendoza et al., 2018).

## Results

### QPCR Primers and Probes

To select primer/probe sets that maximize detection of HIV-1 we analyzed four previously characterized candidates *in silico* using intact proviral genomes from the Los Alamos HIV sequence database (Bruner and Siliciano, 2018; Bruner et al., 2019; Palmer et al., 2003; Schmid et al., 2010). The selected primer/probes cover conserved regions in the HIV-1 genome including the packaging signal (*PS*), group-specific antigen (*gag*), polymerase (*pol*), and envelope (*env*) (Fig. S1). Allowing for 1 and up to 4 mismatches (3 mismatches at the 5’ end and 1 mismatch at the 3’ end) in probes and primers respectively the *PS, gag, pol* and *env* primer/probes detected 72%, 83%, 94% and 92% of 578 intact clade B sequences in the Los Alamos HIV sequence database (Fig. 1A and Fig. S2) (Lefever et al., 2013; Rutsaert et al., 2018; Stadhouders et al., 2010). All genomes scored positive with at least one of the four primer/probe sets. Notably, the large majority (99%) of genomes were positive for at least one of the many combinations of two primer/probe sets. However, any single 2 probe combination was at best 86% sensitive (*pol+env*).

**Fig. 1.**
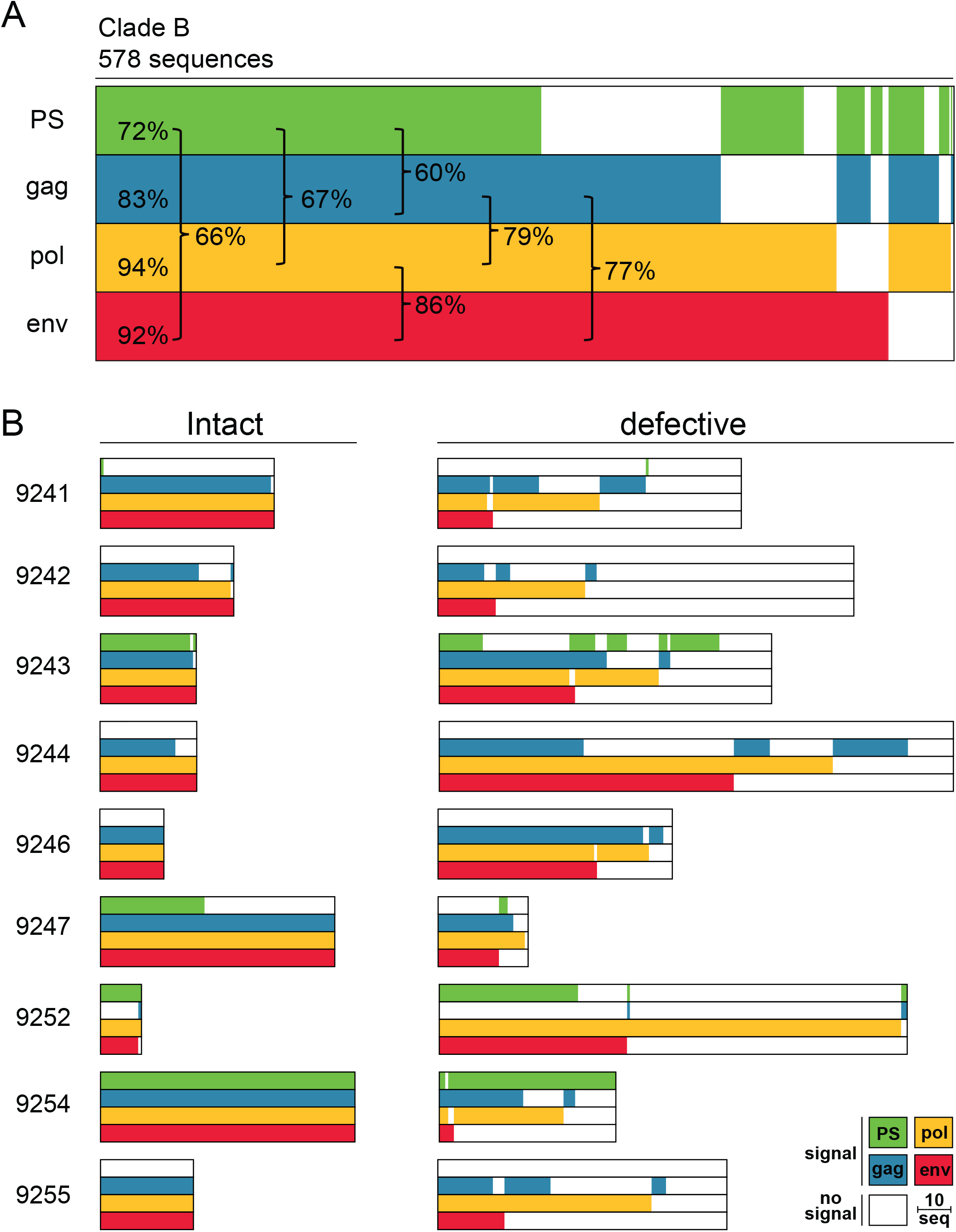
Predicted detection of HIV-1. (A) Horizontal bars represent the predicted detection of 578 intact clade B proviral sequences (Los Alamos HIV sequence database) by qPCR primer/probe sets that target *PS* (green), *gag* (blue), *pol* (yellow) and *env* (red) regions. Signal prediction for each individual proviral sequence is represented by the presence of the color of the respective primer/probe set. Sequences containing polymorphisms that prevent signal detection are represented by the absence of color. The percentage indicates the fraction of detected sequences for individual primer/probe sets or combinations of two primer/probe sets (brackets). (B) Horizontal bars represent the predicted detection of 401 intact and 977 defective near-full length genomes from 9 individuals (Lu et al., 2018; Mendoza et al., 2018). The same primer/probe sets and color scheme is used as described above. The group of defective sequences includes near-full length genomes that carry small insertions, deletions and defects in the packaging site and/or major splice donor. The length of the scale bar represents 10 proviral sequences.

To test whether these primer/probe sets can discriminate between intact and defective proviruses we also performed the same *in silico* analysis on 1378 intact and defective HIV-1 sequences from 9 individuals that received a combination of two broadly neutralizing monoclonal antibodies during treatment interruption (Lu et al., 2018). In six out of nine patients we observed HIV-1 sequence polymorphisms that cause a predicted loss of signal for at least one of the primer/probe sets (Fig. 1B). For example, intact viruses in individuals 9241, 9242, 9244, 9246 and 9255 are predicted to be negative for the *PS* primer/probe set. In addition to the problem of sensitivity, 2 probe combinations also have a potential problem with specificity since a number of defective viruses were predicted to be positive for several of the 2 probe combinations tested. The potential magnitude of this problem varies with the probe combination and the individual analyzed. For example, in 9252 80% of the viruses detected with the *PS*+*env* combination are defective, whereas in 9243 it’s 35% (Fig 1B). Thus, the *in silico* data suggest that any single combination of 2 probes would not be sufficient for sensitive and specific reservoir measurements due to HIV-1 sequence polymorphisms within and between individuals.

### Quadruplex qPCR (Q4PCR)

To accommodate HIV-1 sequence diversity we developed a multiplex qPCR strategy for simultaneous detection of 4 probes, *PS, gag, pol*, and *env* using a 384-well format (Q4PCR). Using this approach we analyzed samples from 2 separate time points from 6 individuals enrolled in a clinical trial that involved analytical treatment interruption after infusion of a combination of two broadly neutralizing monoclonal antibodies (Lu et al., 2018; Mendoza et al., 2018).

Proviral genomes were amplified from DNA extracted from purified CD4^+^ T cells obtained 2 weeks before (wk-2) and 12 weeks (wk12) after treatment interruption. To determine overall HIV-1 proviral frequency, genomic DNA from CD4^+^ T cells was assayed for *gag* by qPCR. We found *gag*^+^ proviruses with the expected variation between individuals at a median frequency of 417 out of 10^6^ CD4^+^ T cells (Table S1). DNA from CD4^+^ T cells was diluted to a concentration equivalent to single copy of *gag* per reaction and assayed by Q4PCR (Fig. 2).

**Fig. 2.**
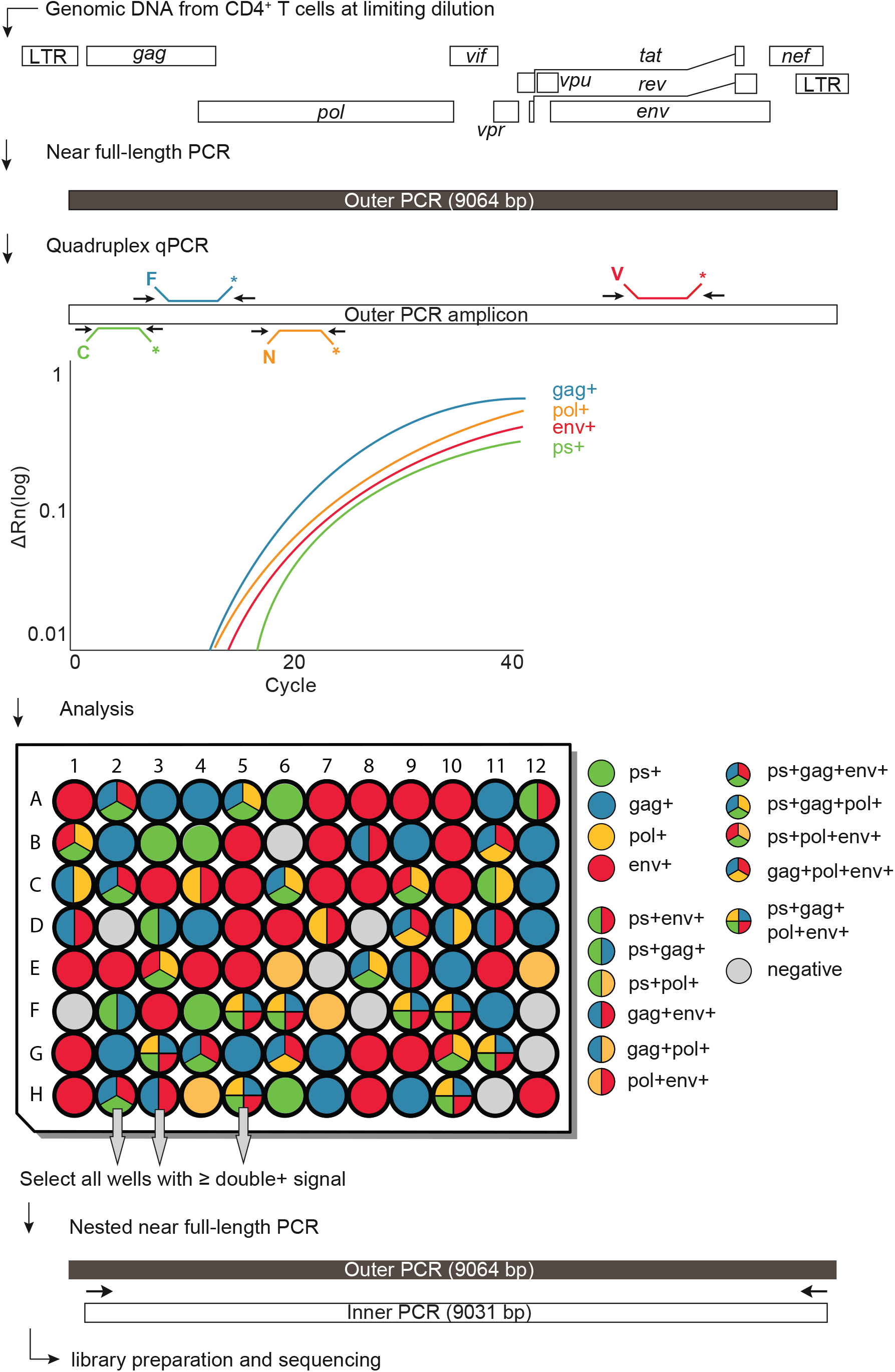
Quadruplex qPCR approach (Q4PCR) Schematic representation of the Q4PCR protocol. Genomic DNA from CD4^+^ T cells was subjected to limiting dilution qPCR with a *gag* specific primer/probe set to determine overall HIV-1 proviral frequency. Near full-length proviral genomes were amplified from CD4^+^ T cell genomic DNA in samples diluted to single copy concentrations based on *gag* qPCR. An aliquot of the resulting amplicons was assayed by quadruplex qPCR using a combination of primer/probe sets covering *PS, gag, pol* and *en*v. Samples with positive signal for any combination of at least two primer/probe sets were collected and subjected to nested near full-length PCR, library preparation and next generation sequencing.

Individual reactions containing a single proviral copy were amplified to produce near full-length and subgenomic proviruses using 5’ of *gag* and LTR primers (Ho et al., 2013; Li et al., 2007). Individual amplicons were then tested for reactivity with each of the 4 qPCR probes. Participant 9254 was excluded from the quantitative analysis because of inadequate sample availability.

### Quantitative Analysis

The number of samples showing reactivity with any one of the selected HIV-1 probes after near full-length amplification was similar (1.5-fold lower) to the number predicted from short segment *gag* qPCR performed before the amplification indicating that near full-length genome PCR reaction was efficient (Fig. 3A and Table S1).

**Fig. 3.**
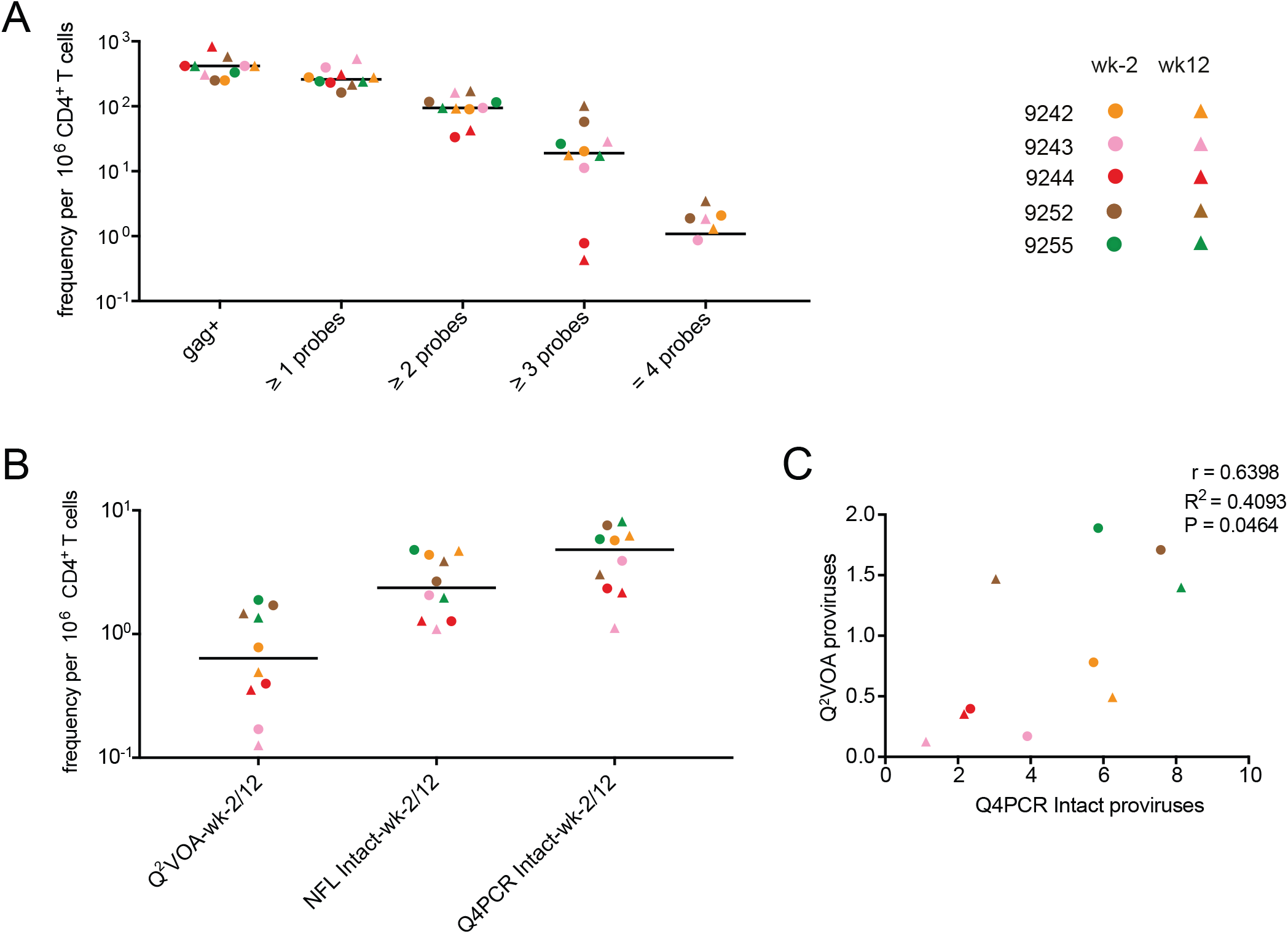
Quantitative analysis. (A) Frequency per million CD4^+^ T cells of: *gag*^+^ proviruses amplified from genomic DNA and samples with any 1, or 2, or 3 or all 4 qPCR probe signals after near full-length amplification for the preinfusion (wk-2) and week 12 (wk12) time points. Horizontal bars indicate median values. For patient 9242 the frequency of *env*^*+*^ proviruses amplified from genomic DNA per million CD4^+^ T cells is plotted due to limited *gag*^*+*^ amplification signal. (B) Comparison of frequencies of inducible proviruses (Q^2^VOA), intact proviruses obtained with NFL sequencing strategy (NFL Intact) and intact proviruses identified with Q4PCR (Q4PCR Intact) at preinfusion and week 12 time points for the same samples (Lu et al., 2018; Mendoza et al., 2018). (C) Pearson correlation between frequency of intact proviruses identified with Q4PCR and inducible proviruses measured by Q^2^VOA at preinfusion and week 12 time points (Lu et al., 2018; Mendoza et al., 2018). Participant 9254 was excluded from the quantitative analysis because of inadequate sample availability. Individual patients are depicted in different colors. Time points are represented by circles and triangles for week −2 and week 12 respectively.

There was substantial variation between individuals (see below), but the median frequency of proviruses reactive to 2 or more probes was 94 × 10^-6^ CD4^+^ T cells or approximately 4x less than *gag* DNA alone. The frequency of proviruses that scored positive with at least 3 or all 4 probes was lower still at 19 × 10^-6^ and 1 × 10^-6^ CD4^+^ T cells (Fig. 3A and Table S1).

To determine which of the proviruses detected by Q4PCR were intact we performed next generation sequencing (Fig. 2). Preliminary sequence analysis indicated that samples reacting with only a single primer/probe or only the *PS+gag* combination were defective, and these samples were mostly omitted from further analysis. All other samples showing reactivity with 2 or more of the 4 qPCR probes were sequenced.

1832 proviruses were sequenced. On average we obtained 153 proviral sequences per time point per participant. Proviral sequences were scored as intact if they did not have deletions or insertions, were in frame, did not contain stop codons, and had intact packaging signals and major splice donors (MSDs) (Fig. S3). In total we found 237 intact and 1595 defective proviral sequences. Intact proviruses were found at a median frequency of 4.8X10^-6^ CD4^+^ T cells, a nearly 20-fold lower frequency than proviruses reacting with 2 or more qPCR probes (Fig 3A and B). NFL sequencing and Q^2^VOA performed on the same samples showed 2-fold and 7.6-fold fewer intact and inducible proviruses per million CD4^+^ T cells than Q4PCR respectively (Lu et al., 2018; Mendoza et al., 2018) (Fig. 3B and Table S2). Despite the limited sample size, there was a significant correlation between the number of viruses measured by outgrowth assays and Q4PCR (Fig 3C). Furthermore, the detection of intact proviruses by Q4PCR was highly reproducible. In two sets of independent experiments performed on samples from 4 individuals there was a strong agreement (Pearson r=0.7764, P=0.0235) consistent with 1.5-fold variability in the assay (Fig. S4).

### Comparison with Q^2^VOA and NFL Sequencing

To compare the intact proviruses obtained by Q4PCR with those obtained by Q^2^VOA and NFL sequencing we created Euler diagrams and phylogenetic trees using *env* sequences (Lu et al., 2018; Mendoza et al., 2018) (Fig. 4A, Fig. S5 and Table S3). There was a substantial overlap between *env* sequences identified by all three methods. Among all intact Q4PCR sequences 68% and 66% were identical to intact NFL and Q^2^VOA sequences, respectively. Thus, we have been unable to detect a selection bias in the HIV-1 reservoir sequences obtained by Q4PCR compared to Q^2^VOA or NFL sequencing. In line with our previous observations from both Q^2^VOA and NFL sequencing, the new Q4PCR sequencing strategy was unable to detect a sequence match between intact proviral sequences and rebound viruses obtained by single genome analysis (SGA) at the time of rebound (Lu et al., 2018).

**Fig. 4.**
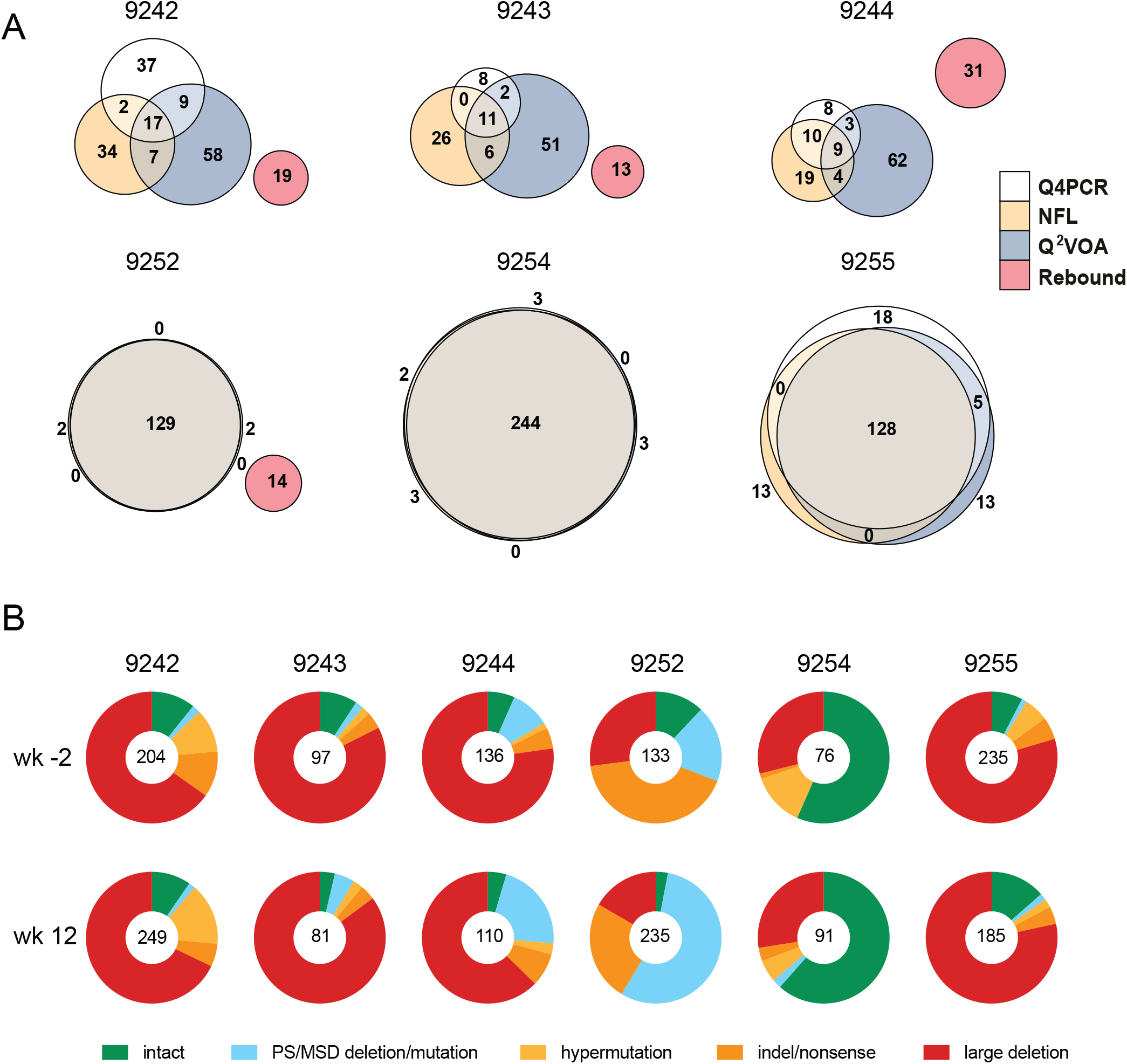
Qualitative sequence analysis. (A) Euler diagrams representing the overlap between *env* sequences obtained from Q4PCR (white), Q^2^VOA (blue), NFL sequencing (yellow) and rebound plasma SGA or PBMC outgrowth culture (red) from participants 9242, 9243, 9244, 9252, 9254 and 9255. (Lu et al., 2018; Mendoza et al., 2018). Q4PCR, Q^2^VOA and NFL sequences obtained from the preinfusion and week 12 time point were combined. Identical *env* sequences were considered as shared sequences. The number inside overlapping areas is the sum of all shared sequences. (B) Pie charts depict the distribution of intact and defective proviral sequences at the preinfusion (wk-2) and week 12 (wk12) time points. The number in the middle of the pie represents the total number of proviruses sequenced. Pie slices indicate the proportion of sequences that were intact or had different defects, including packaging signal defects and MSD site mutations (blue), premature stop codons mediated by hypermutation (yellow), single nucleotide indels or nonsense mutations (orange) and sequences with large internal deletions (red).

### Qualitative Analysis

To determine how integrated HIV-1 varied between individuals and time points within an individual we analyzed both defective and intact proviruses (Fig. 4B and Table S4). Defective viruses were further divided into those with defective packaging signals or MSDs, hypermutation, indels or nonsense mutations, and large deletions. The distribution of defective and intact viruses was relatively constant between the 2 time points in each individual with the exception of 9252 who showed a relative decrease in the number of intact proviruses and an increased representation of proviruses with defective packaging signals or MSDs between weeks −2 and 12. Overall, large internal deletions were the most frequent source of defective proviruses followed by packaging site and/or MSD defects, indel/nonsense mutations and hypermutation. However, the contribution of intact proviruses and individual categories of defects varied significantly between individuals. For example, when the 2 time points are combined the fraction of intact proviruses in individual 9244 was only 5.7% compared to 59% for 9254.

### Sensitivity and Predictive Value

To examine individual probes and probe combinations for their ability to identify intact proviruses, we compared sequencing results with qPCR and determined the predictive value of each probe. The positive predictive value (PPV) is the probability that a sample that is reactive with a specific probe or combination of probes is associated with an intact proviral sequence.

Of a total of 1832 samples assayed, the majority were positive for only 2 (n=908) followed by 3 (n=615), 1 (n=270) and 4 (n=39) probe signals (Fig. 5A). Notably, no sample with an isolated single probe signal was associated with an intact proviral sequence, and only 6% of all samples that were positive for any combination of 2 probes contained intact proviruses. In contrast, 26% and 51% of all samples that were reactive with any combination of 3 or 4 probes respectively contained intact proviruses. Thus, the positive predictive value is directly correlated to the number of positive probes (Fig. 5B and 5C).

**Fig. 5.**
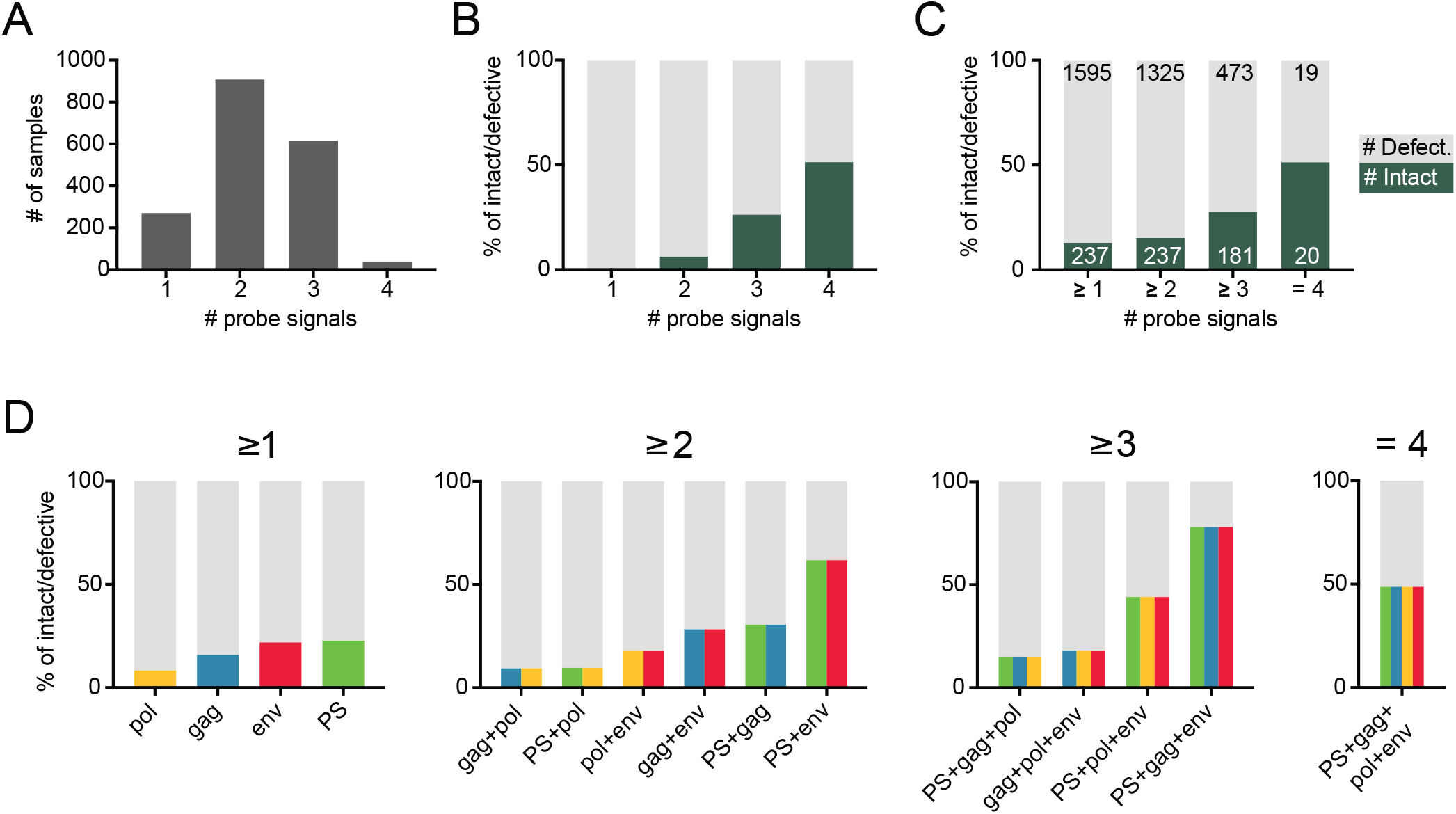
Probe analysis. (A) Bar graphs showing the total number of sequenced samples that scored positive for any 1, or 2, or 3 or all 4 qPCR probes out of a total of 1832 assayed. (B) Stacked bar graphs showing the predictive value for intact (dark green) and defective (grey) proviral genomes of sequenced samples positive for any 1, or 2, or 3 or 4 qPCR probes respectively. (C) Stacked bar graphs showing the predictive value for intact (dark green) and defective (light grey) proviral genomes of sequenced samples positive for at least 1, or 2, or 3 and 4 qPCR probes respectively. The number of samples is depicted in white (intact) or black (defective) respectively. (D) Graphs showing the predictive value for intact and defective proviruses of individual probes and all possible combinations of at least 2, 3 or all 4 probes respectively. The predictive value for intact proviruses is colored for each individual probe *(PS* (green), *gag* (blue), *pol* (yellow) and *env* (red)) or as color combinations for specific probe combinations. The defective fraction is shown in grey.

To determine the contribution of combinations of individual probes, we determined the sensitivity and PPV for all possible probe combinations. Notably, whereas an isolated *env* signal fails to predict intact proviruses, *env* in combination with any other probe shows a very high sensitivity for the detection of intact proviruses, 98% of 237 intact sequences were *env*^*+*^. In contrast, the sensitivity of either *pol*, or *PS* or *gag* with any other probe was lower, 39%, 65% and 83% respectively (Table S5).

The combination of *PS*+*env* has the highest, and *gag+pol* the lowest positive predictive value of any 2 probe combinations (62% and 9.4% respectively, Fig 5D and Table S5). However, even the *PS*+*env* combination has a substantial false discovery rate. Overall 38% of *PS*+*env* positive samples are associated with defective proviral sequences, and 2 of the 6 individuals failed to show any signal with the *PS* primer/probe set (Fig. 6 and Table S6). In both of these cases the absence of the signal was predicted and could be explained by sequence polymorphisms in proviral genomes (Fig.1B and Fig. S6). Thus, polymorphisms limit the applicability of a diagnostic test based on any two probes including *PS*+*env*. Moreover, even in those individuals where the *PS*+*env* combination was effective, the positive predictive value varied from 9.1% to 96% between individuals (Fig. 6 and Table S6).

**Fig. 6.**
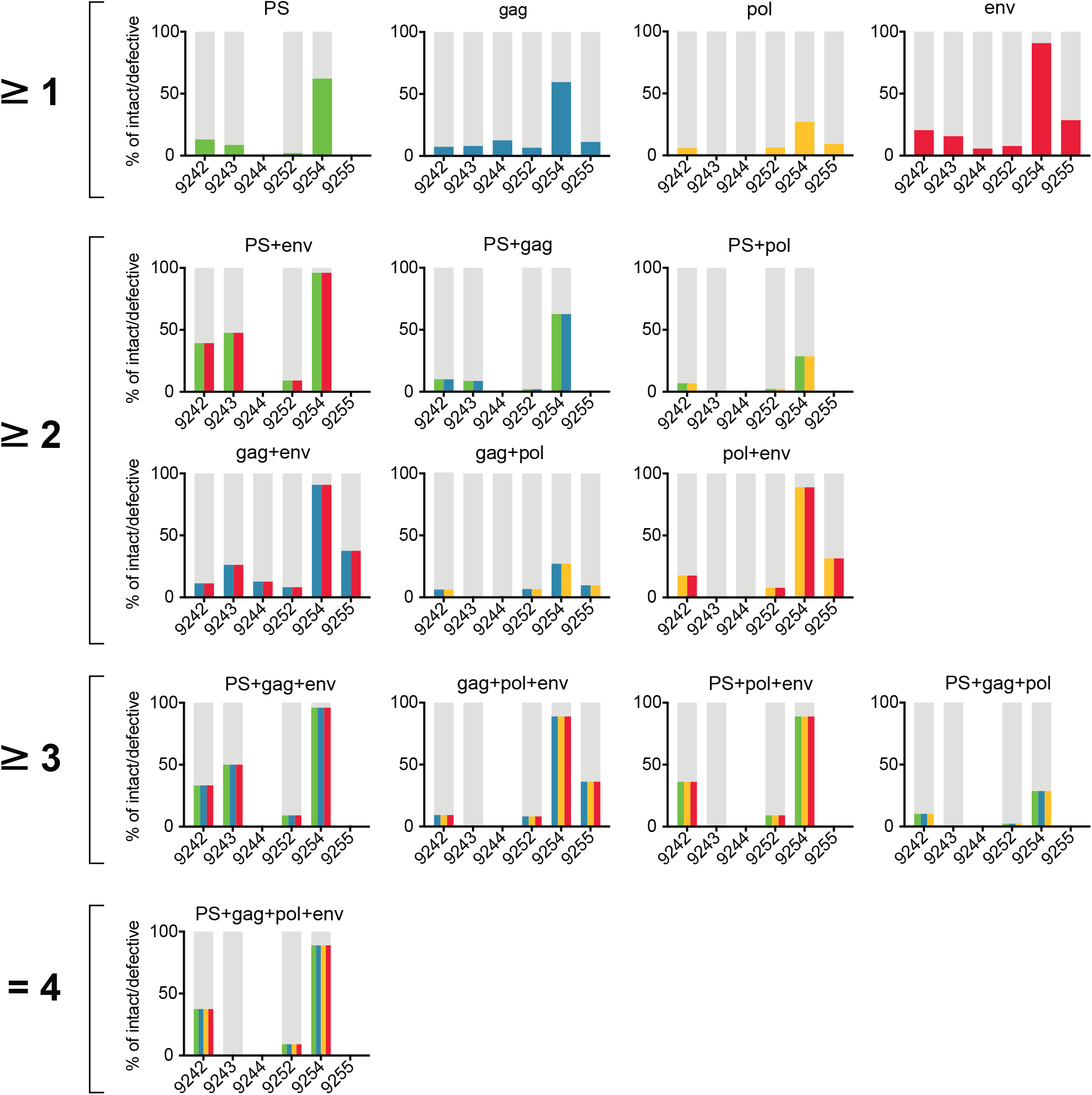
Probe analysis for individual participants. Graphs showing the predictive value for intact and defective proviral genomes of individual probes and all possible combinations of at least 2, 3 or all 4 probes among participants 9242, 9243, 9244, 9252, 9254 and 9255 at the preinfusion (wk-2) and week 12 (wk12) time points. The predictive value for intact proviruses is colored for each individual probe *(PS* (green), *gag* (blue), *pol* (yellow) and *env* (red)) or as color combinations for specific probe combinations. The defective fraction is shown in grey. Blank spaces illustrate the absence of specific probe or probe combination signals in individual patients.

Samples positive for at least 3 probes show considerable improved predictive values. For example, samples that were positive for at least *PS+gag+env* predict intact proviruses at a rate of 78% (Fig 5D). However, the requirement for hybridization with a third or fourth probe decreases the sensitivity of the assay and results in inability to detect 24% and 92% of all positive samples respectively, an effect that appears to be due to HIV-1 sequence variation (Table S5). Only the combination of any two Q4PCR primer/probe sets and next generation sequencing is both sensitive and specific for enumeration of intact proviral genomes.

## Discussion

Accurate measurement of the size of the circulating latent reservoir is essential for evaluating therapeutic interventions that aim to eliminate it (Sengupta and Siliciano, 2018). The combination of Q4PCR and next generation sequencing is a relatively high throughput method that is both sensitive and specific.

Several methods have been used to evaluate the latent HIV-1 reservoir. For example, total integrated proviral DNA can be measured by polymerase chain reaction using *gag* specific primers (Bieniasz et al., 1993; Christopherson et al., 2000; Palmer et al., 2003). This assay is simple and quantitative. However, it does not distinguish between rare replication-competent and more abundant replication-defective proviruses (Bruner et al., 2016; Eriksson et al., 2013; Ho et al., 2013). Thus, the overwhelming background of defective proviruses would make any change in the replication competent latent reservoir difficult to detect using this assay. Viral outgrowth assays measure infectious viruses that can be recovered by CD4^+^ T cell activation *in vitro* (Chun et al., 1997; Lorenzi et al., 2016). However, these assays are very labor intensive, variable and not sensitive to any change below a factor of 6 (Crooks et al., 2015; Rosenbloom et al., 2019). In addition, these assays selectively underestimate the reservoir because only a fraction of latently infected cells can be induced after a single round of stimulation *in vitro* (Ho et al., 2013; Hosmane et al., 2017).

Sequencing near full-length proviral genomes from limiting dilution CD4^+^ T cell DNA samples allows relatively unbiased characterization of the proviral reservoir (Hiener et al., 2017; Ho et al., 2013; Lu et al., 2018). This technique is very labor intensive and requires interrogation of thousands of reactions per sample making these studies challenging. In addition there is some variation between the number of estimated intact genomes measured by DNA sequencing assays based on Sanger and next generation sequencing technologies 12-114 × 10^-6^ (Bruner et al., 2016; Ho et al., 2013) vs. 2.8-24 × 10^-6^ CD4^+^ T cells (Hiener et al., 2017; Lee et al., 2017; Lu et al., 2018; Sharaf et al., 2018; Vibholm et al., 2019). One possible explanation for the observed difference is the use of an empirical Bayesian model to estimate the number of intact proviral genomes in the Sanger-based studies necessitated by the absence of detected intact genomes in 13 out of 26 subjects (Bruner et al., 2016; Ho et al., 2013).

Intact Proviral DNA Assay (IPDA) is a high throughput method to measure integrated proviral DNA using droplet digital PCR to probe for the presence of *PS*+*env* in the proviral genome (Bruner et al., 2019). This method detects intact proviruses at a higher frequency than the sequencing methods (100 × 10^-6^ CD4^+^ T cells) (Bruner et al., 2019). The IPDA relies on amplification of two subgenomic regions that together sample 222bp or only 2% of the 9.7kb HIV-1 genome. As a result, a significant fraction of proviruses are incorrectly categorized as intact which leads to an overestimation of intact proviral DNA (Bruner et al., 2019). Moreover, our experiments indicate that polymorphisms lead to variation in the levels of sensitivity and specificity of the *PS*+*env* probe combination in different individuals suggesting that any single combination of 2 probes would not be sufficient for broadly applicable reservoir measurements. Most importantly, verification of intact proviruses is not possible in the IPDA and therefore the accuracy of this method will vary between individuals depending on the molecular composition of the reservoir. Nevertheless, the IPDA is a high throughput assay that is more accurate than the total proviral DNA assay, making it the most desirable currently available assay for large studies (Bruner et al., 2019).

The advantage of combining multicolor qPCR and sequencing is that the method is relatively rapid, scalable, and both sensitive and specific. All intact viruses in the 6 individuals analyzed and 99% of the intact Clade B viral sequences in the Los Alamos data base were positive for any combination of 2 of the 4 probes in the Q4PCR reaction. Bar coding and next generation sequencing facilitates the analysis of large numbers of samples simultaneously and enables definitive identification of intact proviruses. In addition, the sequence data provides information on the clonal structure of the latent reservoir and on the nature of the defective proviruses.

In conclusion the combination of 4 probe qPCR and next generation sequencing is a highly sensitive and specific method for measuring intact proviruses in the HIV-1 latent reservoir.

## Materials and methods

### Study Participants

HIV-1-infected participants were enrolled at the Rockefeller University Hospital, New York, USA, and the University Hospital Cologne, Cologne, Germany, in an open-label phase 1b study (http://www.clinicaltrials.gov; NCT02825797; EudraCT: 2016-002803-25) (Mendoza et al., 2018). All participants provided written informed consent before participation in the study and the study was conducted in accordance with Good Clinical Practice. The protocol was approved by the Federal Drug Administration in the USA, the Paul-Ehrlich-Institute in Germany, and the Institutional Review Boards (IRBs) at the Rockefeller University and the University of Cologne. Participants received the combination of two broadly neutralizing antibodies (3BNC117 and 10-1074) intravenously at a dose of 30 mg kg^-1^ body weight of each antibody, at weeks 0, 3 and 6, unless viral rebound occurred. ART was discontinued 2 days after the first infusion of antibodies (day 2). Leukapheresis was performed at the Rockefeller University Hospital or at the University Hospital Cologne at week –2 and week 12. Peripheral blood mononuclear cells (PBMCs) were isolated by density gradient centrifugation and cells were cryopreserved in fetal bovine serum plus 10% DMSO.

### Bioinformatic binding prediction of qPCR primers and probes

Previously characterized primers and probes were mapped to HXB2 to identify regions of binding (Althaus et al., 2010; Bruner and Siliciano, 2018; Bruner et al., 2019; Malnati et al., 2008; Palmer et al., 2003; Schmid et al., 2010). Then, we selected all HIV-1 Clade B (n=578), Clade C (n=456) and Clade A (n=94) full-length intact proviral genomes from the Los Alamos HIV sequence database (https://www.hiv.lanl.gov/content/sequence/NEWALIGN/align.html) that included sequence information for the packaging signal. After alignment to the HXB2 reference sequence using MAFFT v.7.309, we predicted binding and amplification signal for primer/probe sets with a maximum of one probe mismatch and four primer mismatches. Forward and reverse primers were further subdivided in a 5’ and 3’ end and a maximum of three mismatches at the 5’ end and one mismatch at the 3’ end was allowed for predicted binding and positive signal (Lefever et al., 2013; Rutsaert et al., 2018; Stadhouders et al., 2010). In addition, we used Geneious 11 and Primer3 to assess physical properties such as melting temperatures and secondary structures. Based on predicted binding and favorable physical properties for multiplex qPCR reactions we selected the packaging signal (*PS*) (Bruner and Siliciano, 2018), group-specific antigen (*gag*) (Palmer et al., 2003), polymerase (*pol*) (Schmid et al., 2010), and envelope (*env*) (Bruner et al., 2019) primer/probe sets. To test whether these primer/probe sets can discriminate between intact and defective proviruses we analyzed 1378 intact and defective near full-length HIV-1 sequences from 9 individuals (Lu et al., 2018).

### CD4^+^ T cell Isolation

Total CD4^+^ T cells were isolated from cryopreserved PBMCs by manual magnetic labeling and negative selection using the CD4^+^ T Cell Isolation Kit (Miltenyi Biotec).

### DNA Isolation and Quantification

Genomic DNA from 1-10 Million CD4^+^ T cells was isolated using the Gentra Puregene Cell Kit (Qiagen). In some experiments, DNA was isolated using phenol-chloroform (Klein et al., 2011). Briefly, CD4^+^ T cells were lysed in Proteinase K buffer [100 mM Tris (pH 8), 0.2% SDS, 200 mM NaCl, 5 mM EDTA] and 20 mg/mL Proteinase K at 56 °C for 12 hours followed by genomic DNA extraction with phenol/chloroform/isoamyl alcohol extraction and ethanol precipitation. The Qubit 3.0 Fluorometer and Qubit dsDNA BR Assay Kit (Thermo Fisher Scientific) was used to measure DNA concentrations.

### Limiting Dilution *gag* quantitative PCR

Genomic DNA was assayed in a 384-well plate format using the Applied Biosystem QuantStudio 6 Flex Real-Time PCR System. HIV-1-specific primers and a probe targeting a conserved region in *gag* were used in a limiting dilution quantitative PCR (qPCR) reaction (Forward primer 5’-ATGTTTTCAGCATTATCAGAAGGA-3’, Internal probe 5’-/6-FAM/CCACCCCAC/ZEN/AAGATTTAAACACCATGCTAA/3’/IABkFQ/, Reverse primer 5’-TGCTTGATGTCCCCCCACT-3’ (Integrated DNA Technologies) (Palmer et al., 2003).

Each qPCR reaction was carried out in a 10 µl total reaction volume containing 5µl of TaqMan Universal PCR Master Mix containing Rox (Applied Biosystems, Catalog no. 4304437), 1 µl of diluted genomic DNA, nuclease free water and the following primer and probe concentrations: 337,5nM of Forward and Reverse primers with 93,75nm of *gag* internal probe. *Gag* qPCR conditions were 94°C for 10 min; 50 cycles of 94°C for 15 s and 60 °C for 60 s.

Genomic DNA was serially diluted to concentrations ranging from 2000 to 250 CD4^+^ T cells per µl with a minimum of 24 reactions per concentration. We selected DNA dilutions wherein <30% of the *gag* PCR reactions were positive for further analysis because they have more than an 80% probability of containing a single copy of HIV-1 DNA in each PCR reaction based on the Poisson distribution (Lu et al., 2018).

### Near Full-Length HIV-1 PCR (1.PCR)

We used a two-step nested PCR approach to amplify near full-length HIV-1 genomes. All reactions were carried out in a 20 µl reaction volume using Platinum Taq High Fidelity polymerase (Thermo Fisher Scientific). The outer PCR reaction was performed on genomic DNA at the previously determined single copy dilution using outer PCR primers BLOuterF (5’-AAATCTCTAGCAGTGGCGCCCGAACAG-3’) and BLOuterR (5’ - TGAGGGATCTCTAGTTACCAGAGTC - 3’). Touchdown cycling conditions were: 94°C for 2 m; then 94°C for 30 s, 64°C for 30 s, 68°C for 10 m for 3 cycles; 94°C for 30s, 61°C for 30 s, 68°C for 10 m for 3 cycles; 94°C for 30 s, 58°C for 30 s, 68°C for 10 m for 3 cycles; 94°C for 30 s, 55°C for 30 s, 68°C for 10 m for 41 cycles; then 68°C for 10 m (Ho et al., 2013; Li et al., 2007).

### Quadruplex qPCR

Undiluted 1 µl aliquots of the near full-length 1.PCR product were subjected to a quadruplex qPCR reaction using a combination of four primer/probe sets that target conserved regions in the HIV-1 genome. Each primer/probe set consists of a forward and reverse primer pair as well as a fluorescently labeled internal hydrolysis probe: **Packaging Signal *(PS)*** (Forward: 5’-TCTCTCGACGCAGGACTC-3’, Reverse: 5’-TCTAGCCTCCGCTAGTCAAA-3’,Probe: 5’/Cy5/TTTGGCGTA/TAO/CTCACCAGTCGCC/3’/IAbRQSp) (Integrated DNA Technologies) (Bruner and Siliciano, 2018), ***env*** (Forward: 5’-AGTGGTGCAGAGAGAAAAAAGAGC-3’, Reverse: 5’-GTCTGGCCTGTACCGTCAGC-3’, Probe: 5’/VIC/CCTTGGGTTCTTGGGA/3’/MGB) (Thermo Fisher Scientific) (Bruner et al., 2019), ***gag*** (Forward: 5’-ATGTTTTCAGCATTATCAGAAGGA-3’, Reverse: 5’-TGCTTGATGTCCCCCCACT-3’, Probe: 5’/6-FAM/CCACCCCAC/ZEN/AAGATTTAAACACCATGCTAA/3’/IABkFQ) (Integrated DNA Technologies) (Palmer et al., 2003) and ***pol*** (Forward: 5’-GCACTTTAAATTTTCCCATTAGTCCTA-3’, Reverse: 5’-CAAATTTCTACTAATGCTTTTATTTTTTC-3’, Probe: 5’/NED/AAGCCAGGAATGGATGGCC/3’/MGB) (Thermo Fisher Scientific) (Schmid et al., 2010).

Each quadruplex qPCR reaction was carried out in a 10 µl total reaction volume containing 5µl of TaqMan Universal PCR Master Mix containing Rox (Applied Biosystems, Catalog no. 4304437), 1 µl of diluted genomic DNA, nuclease free water and the following primer and probe concentrations: **Packaging Signal *(PS)*** 675 nM of Forward and Reverse primers with 187.5 nM of *PS* internal probe, ***env*** 90 nM of Forward and Reverse primers with 25 nM of *env* internal probe, ***gag*** 337.5nM of Forward and Reverse primers with 93.75 nM of *gag* internal probe, ***pol*** 675 nM of Forward and Reverse primers with 187.5 nM of *pol* internal probe. qPCR conditions were 94°C for 10 min; 40 cycles of 94 °C for 15 s and 60 °C for 60s. All qPCR reactions were performed in a 384-well plate format using the Applied Biosystem QuantStudio 6 Flex Real-Time PCR System.

### qPCR data analysis

We used QuantStudio™ Real-Time PCR Software, Version 1.3 (Thermo Fisher Scientific), for data analysis. The same baseline correction (Start Cycle: 3, End Cycle: 10) and normalized reporter signal (ΔRn) threshold (ΔRn threshold=0.025) was set manually for all targets/probes. Fluorescent signal above the threshold was used to determine the threshold cycle (Ct).

Samples with a Ct value between 10 and 40 of any probe or probe combination were identified. Preliminary sequence analysis indicated that samples reacting with only a single primer/probe or only the *PS+gag* combination were defective, and these samples were mostly omitted from further analysis. All other samples showing reactivity with two or more of the four qPCR probes were selected for further processing

### Nested Near Full-Length HIV-1 PCR

The nested PCR reaction was performed on undiluted 1 µl aliquots of the near full-length 1.PCR product. Reactions were carried out in a 20 µl reaction volume using Platinum Taq High Fidelity polymerase (Thermo Fisher Scientific) and PCR primers 275F (5’-ACAGGGACCTGAAAGCGAAAG-3’) and 280R (5’ - CTAGTTACCAGAGTCACACAACAGACG-3’) (Ho et al., 2013) at a concentration of 800nM. Touchdown cycling conditions were: 94°C for 2 m; then 94°C for 30 s, 64°C for 30 s, 68°C for 10 m for 3 cycles; 94°C for 30s, 61°C for 30 s, 68°C for 10 m for 3 cycles; 94°C for 30 s, 58°C for 30 s, 68°C for 10 m for 3 cycles; 94°C for 30 s, 55°C for 30 s, 68°C for 10 m for 41 cycles; then 68°C for 10 m.

### Library preparation and sequencing

All nested PCR products were subjected to library preparation without prior gel visualization. The Qubit 3.0 Fluorometer and Qubit dsDNA BR Assay Kit (Thermo Fisher Scientific) was used to measure DNA concentrations. Samples were diluted to a concentration of 10-20ng/µl. Tagmentation reactions were carried out using 1µl of diluted DNA, 0,25µl Nextera TDE1 enzyme and 1,25µl Nextera TD buffer (Illumina). Tagmented DNA was ligated to unique i5/i7 barcoded primer combinations using the Illumina Nextera XT Index Kit v2 and KAPA HiFi HotStart ReadyMix (2X) (KAPA Biosystems), and then purified using AmPure Beads XP (Agencourt). Three hundred eighty-four purified samples were pooled into one library and then subjected to paired-end sequencing using Illumina MiSeq Nano 300 V2 cycle kits (Illumina) at a concentration of 12 pM.

### Sequence Assembly

HIV sequence reconstruction was performed by our in-house pipeline called HIVA (HIV Assembler), a pipeline for the assembly of raw sequencing reads into annotated HIV genomes, capable of reconstructing thousands of genomes within hours. First, a quality-control check trims Illumina adapters and low-quality bases, followed by multiple assembly steps that combine two widely used classes of algorithms: de-Bruijn-graph (DBG) and overlap-layout-consensus (OLC). DBG is performed by SPAdes for initial de novo assembly of contigs which are aligned via BLAST to a database of HIV genome sequences to select the closest reference. OLC is performed by MIRA in two steps. First, a modified version of the closest reference is generated by alignment to the contigs produced by SPAdes. After, the modified reference is used as a scaffold for the final reference-guided assembly of the initial trimmed reads. Finally, the HIV genome sequence is annotated by alignment to HXB2 using ClustalW. Sequences with double peaks (cutoff consensus identity for any residue <75%) or limited reads (empty wells=<500 sequencing reads) were omitted from downstream analyses. HIVA is constructed with Snakemake, a workflow management system, allowing reproducible data analyses and scalability to cluster and cloud environments.

### Data Availability

All proviral sequences have been deposited in GenBank with the accession codes pending.

### Phylogenetic Analysis

Nucleotide alignments of intact env sequences were translation-aligned using ClustalW v.2.148 under the BLOSUM cost matrix. Sequences with premature stop codons and frameshift mutations that fell in the gp120 surface glycoprotein region were excluded from all analyses. Maximum likelihood phylogenetic trees were then generated from these alignments with RAxML v.8.2.950 under the GTRGAMMA model with 1,000 bootstraps.

### Euler Diagrams

Identical *env* sequences captured by each method (Q^2^VOA, NFL and Q4PRC) were considered as shared sequences. The number inside overlapping areas in the Euler diagram is the sum of all shared sequences. The fraction of shared sequence from individual methods is shown in Table S3.

### Statistical Analyses

Statistical analyses were performed using GraphPad Prism 7.0d for Mac OS X.

## Acknowledgments

We thank all study participants who devoted time to our research; The Rockefeller University Hospital Clinical Research Support Office and nursing staff, the Klein lab and the clinical study group of the Division of Infectious Diseases at the University Hospital Cologne, for help with sample processing; Lillian B. Cohn, Amy S. Huang and all members of the M.C.N. laboratory for helpful discussions and Zoran Jankovic for laboratory support. C. Gaebler is a fellow of the Robert S. Wennett and Mario Cader-Frech Foundation and was supported in part by grant # UL1 TR001866 from the National Center for Advancing Translational Sciences (NCATS, National Institutes of Health (NIH) Clinical and Translational Science Award (CTSA) program), and by the Shapiro-Silverberg Fund for the Advancement of Translational Research. This work was supported by the Bill and Melinda Gates Foundation Collaboration for AIDS Vaccine Discovery Grants OPP1092074, OPP1124068 and OPP1168933; NIH Grants 1UM1 AI100663, and R01AI129795 (to M.C. Nussenzweig); the Einstein–Rockefeller–CUNY Center for AIDS Research (1P30AI124414-01A1); BEAT-HIV Delaney Grant UM1 AI126620 (to M. Caskey); and the Robertson Fund. M.C. Nussenzweig is a Howard Hughes Medical Institute Investigator.

There are patents on 3BNC117 and 10-1074, of which M.C. Nussenzweig is an inventor. M.C. Nussenzweig is a member of the Scientific Advisory Boards of Celldex and Frontier Biotechnologies. The authors declare no additional financial interests.

## Author contributions

C. Gaebler, J.C.C. Lorenzi and M.C. Nussenzweig designed the research. C. Gaebler, J.C.C. Lorenzi, L. Nogueira, C-L. Lu and P. Mendoza performed the research. C. Gaebler, J.C.C. Lorenzi, T.Y. Oliveira, V. Ramos, J. Pai, M. Jankovic, M. Caskey and M.C. Nussenzweig analyzed the data. C. Gaebler, J.C.C. Lorenzi and M.C. Nussenzweig wrote the manuscript.

