## Supplementary material for "Quadruplex qPCR for qualitative and quantitative analysis of the HIV-1 latent reservoir": Figure S1-S6, Table S1-S6

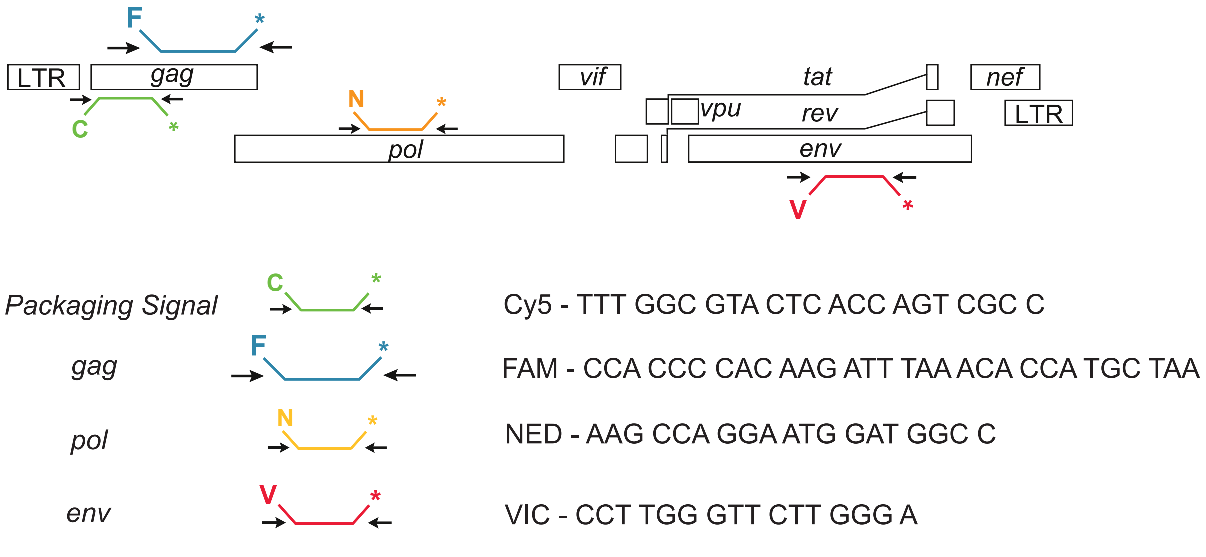


Figure S1. **Q4PCR probes.** Diagram illustrates sequences, fluorophores and HIV-1 genome positions (not at scale) of four probes applied in Q4PCR.

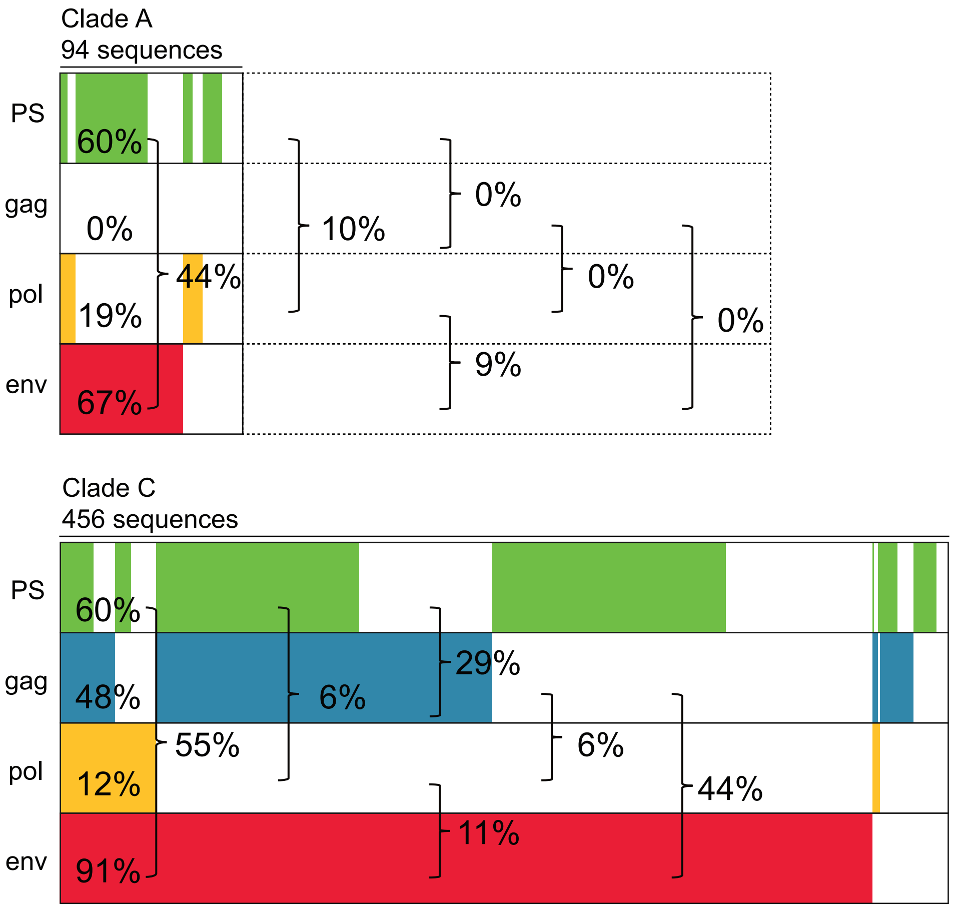


Figure S2. **Predicted Detection of HIV-1 Clade A and C.** Predicted detection of 94 intact clade A and 456 Clade C proviral sequences from the Los Alamos HIV sequence database by primer/probe sets that target *PS* (green), *gag* (blue), *pol* (yellow) and *env* (red) regions. Predicted signals are represented by the presence of the color of the respective primer/probe set. Sequences containing polymorphisms that prevent signal detection are represented by the absence of color. The percentage indicates the fraction of detected sequences for individual primer/probe sets or combinations of two primer/probe sets (brackets).

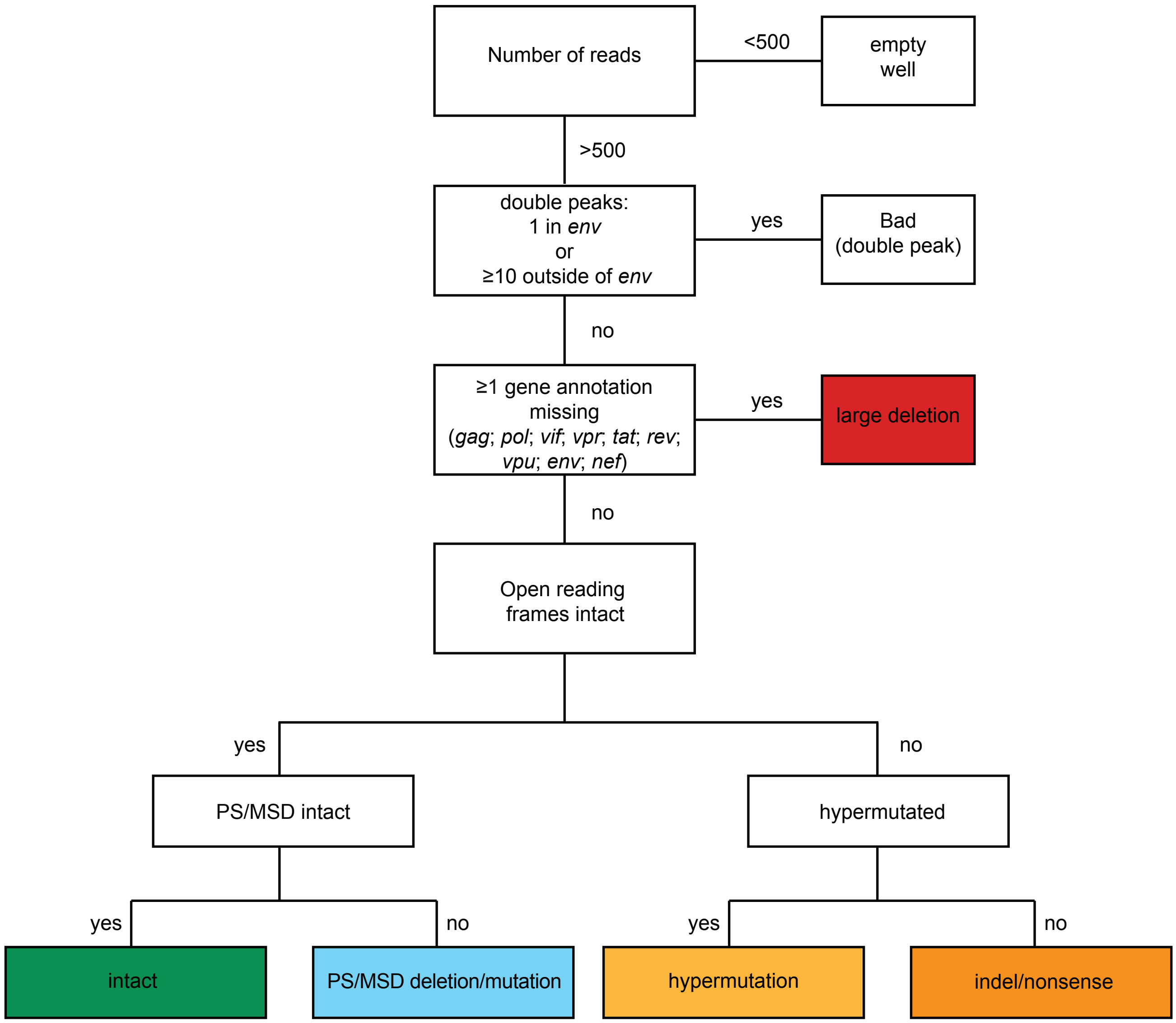


Figure S3. **Sequence classification process.** Sequences with double peaks (cutoff consensus identity for any residue <75%) or limited reads (empty wells=<500 sequencing reads) were omitted from downstream analyses. Assembled HIV genomes were annotated by alignment to HXB2 to identify sequences with missing gene annotations (large deletions), premature stop codons, out-of-frame insertions or deletions (hypermutation or indel), or packaging signal and the major splice donor (MSD) site deletions and mutations.

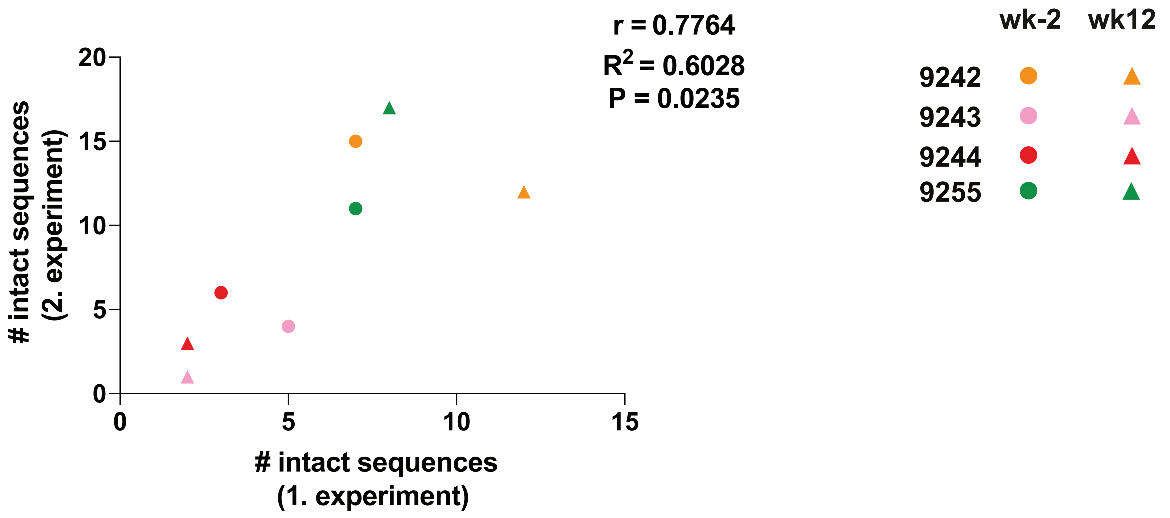


Figure S4. **Assay variability.** Pearson correlation between number of intact proviruses identified in two sets of independent experiments from 4 individuals at preinfusion (wk-2) (circles) and week 12 (triangles) time points.

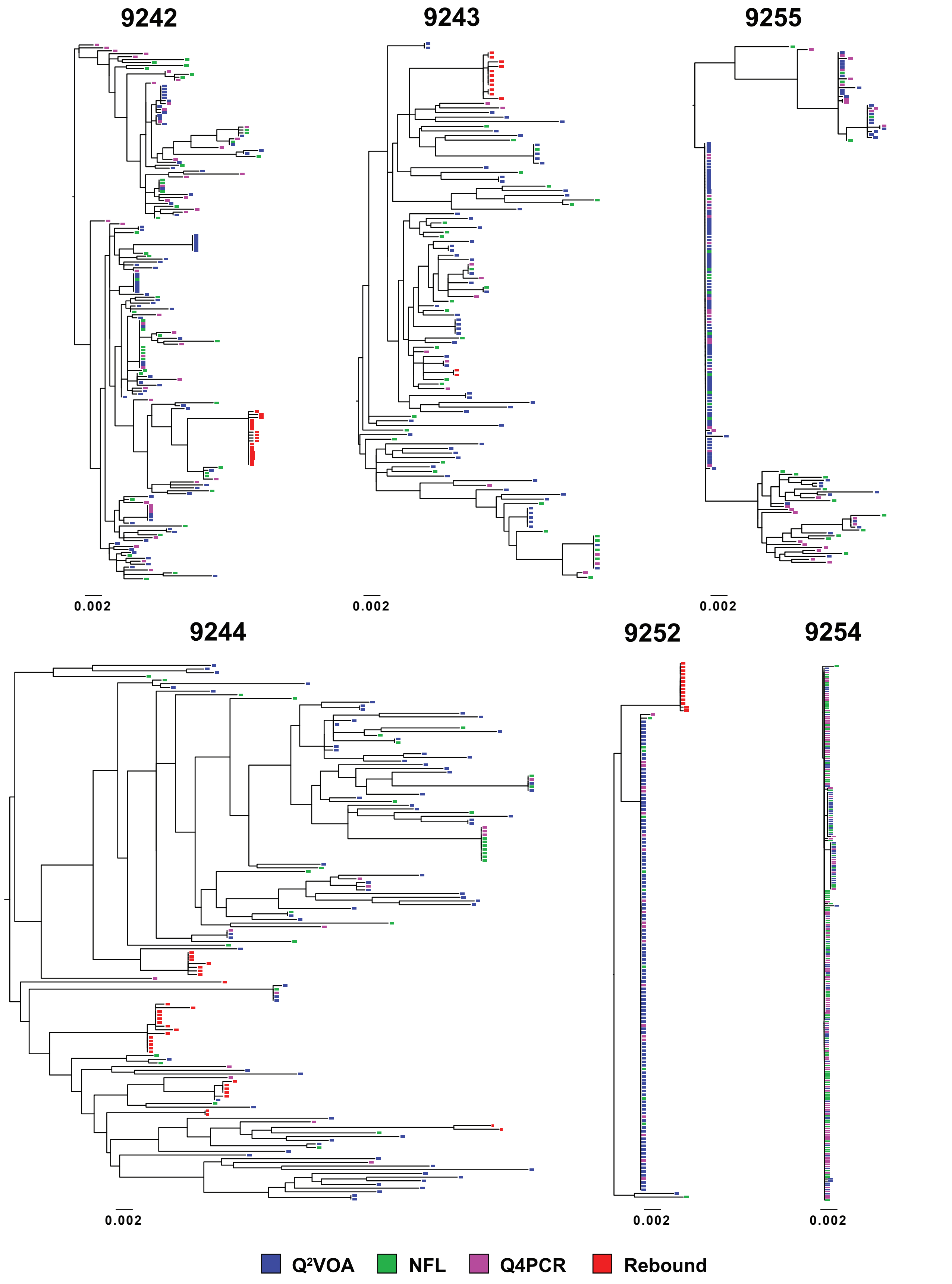


Figure S5. **Phylogenetic trees of env sequences.** Maximum likelihood phylogenetic trees of *env* sequences obtained with Q²VOA (blue), NFL sequencing (green), Q4PCR (purple) and rebound plasma SGA or outgrowth cultures (red).

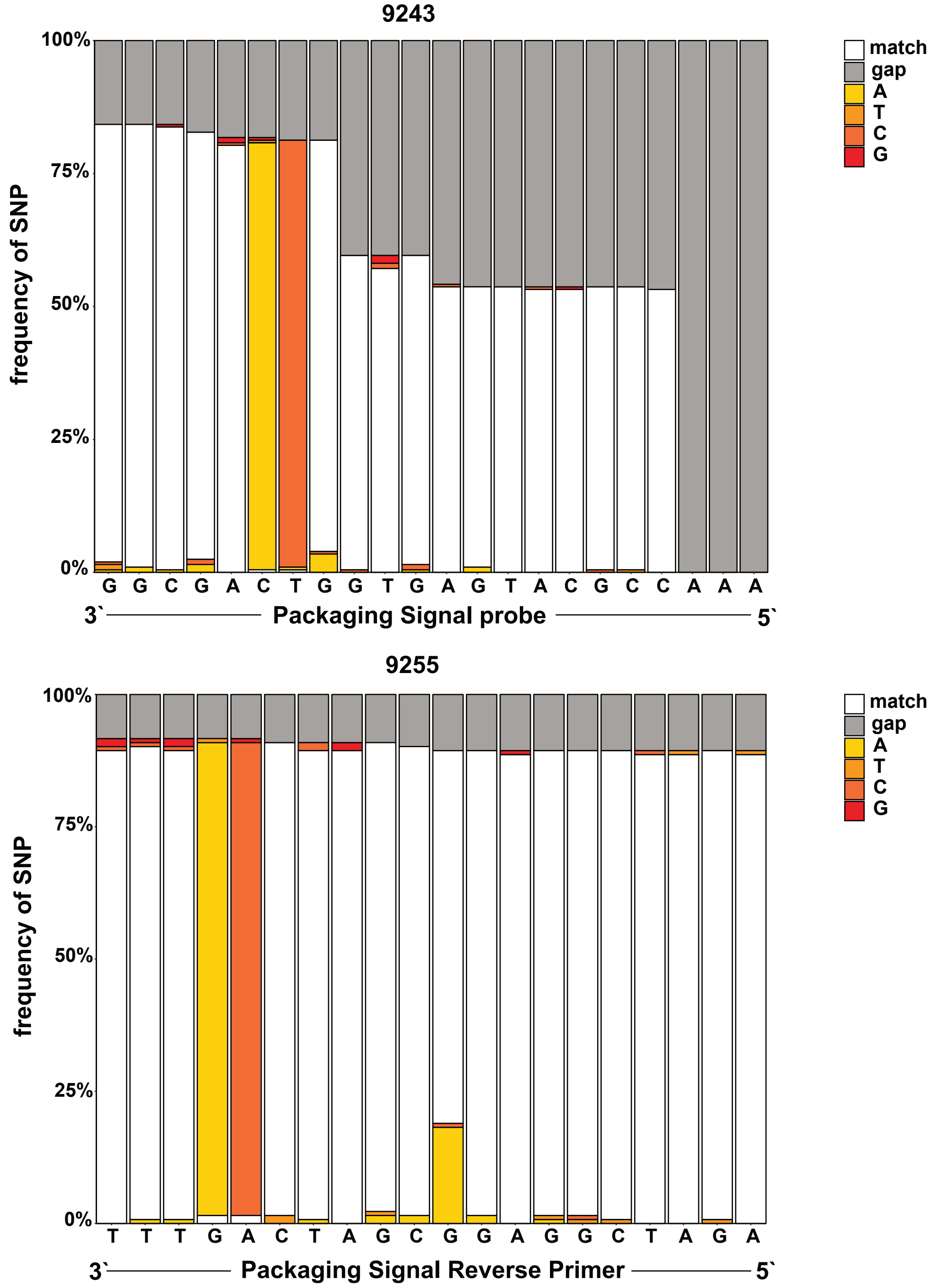


Figure S6. **Single Nucleotide Polymorphism analysis.** Stacked bar graphs depicting sequence identity obtained by aligning all (defective and intact) proviral sequences from patients 9243 and 9255 with *PS* probe and *PS* reverse primer respectively. The frequency of identical nucleotides are shown in white (match). The frequencies of single nucleotide polymorphisms (SNP) that lead to mismatches with tested primer/probe are depicted in yellow (Adenosine), orange (Thymine), light red (Cytosine), dark red (Guanine) and grey (gap in alignment). The *PS* probe and reverse primer are depicted in reverse-complement orientation.

Table S1. **Quantitative analysis of Q4PCR**

| ID | $CD4^+$ T cells tested in Q4PCR | | *Gag*$^+$ (*Env*$^+$*) proviruses per $10^6$ CD4+ T cells | | ≥1 qPCR signal per $10^6$ $CD4^+$ T cells | | ≥2 qPCR signal per $10^6$ $CD4^+$ T cells | | ≥3 qPCR signal per $10^6$ $CD4^+$ T cells | | =4 qPCR signal per $10^6$ $CD4^+$ T cells | |
|---|---|---|---|---|---|---|---|---|---|---|---|---|
| | week -2 | week 12 | week -2 | week 12 | week -2 | week 12 | week -2 | week 12 | week -2 | week 12 | week -2 | week 12 |
| 9242 | 3.84E+06 | 3.84E+06 | 250.0* | 416.7* | 278.6 | 278.1 | 90.1 | 92.7 | 20.3 | 17.7 | 2.1 | 1.3 |
| 9243 | 2.30E+06 | 2.69E+06 | 416.7 | 308.6 | 397.6 | 538.7 | 94.2 | 162.2 | 11.3 | 28.6 | 0.9 | 1.9 |
| 9244 | 3.84E+06 | 2.30E+06 | 416.7 | 833.3 | 231.3 | 309.5 | 33.6 | 43.0 | 0.8 | 0.4 | 0.0 | 0.0 |
| 9252 | 2.11E+06 | 2.30E+06 | 250.0 | 583.3 | 162.4 | 217.0 | 117.0 | 171.9 | 57.8 | 102.0 | 1.9 | 3.5 |
| 9255 | 3.07E+06 | 3.07E+06 | 333.3 | 416.7 | 242.2 | 240.6 | 115.2 | 94.4 | 26.4 | 17.3 | 0.0 | 0.0 |

Table S2. **Quantitative analysis of intact proviruses**

| ID | CD4$^+$ T cells tested in Q$^2$VOA | | CD4$^+$ T cells tested in NFL | | CD4$^+$ T cells tested in Q4PCR | | Inducible proviruses per $10^6$ CD4$^+$ T cells - Q$^2$VOA | | Intact proviruses per $10^6$ CD4$^+$ T cells - NFL | | Intact proviruses per $10^6$ CD4$^+$ T cells - Q4PCR | |
|---|---|---|---|---|---|---|---|---|---|---|---|---|
| | week -2 | week 12 | week -2 | week 12 | week -2 | week 12 | week -2 | week 12 | week -2 | week 12 | week -2 | week 12 |
| 9242 | 7.56E+07 | 1.68E+08 | 4.57E+06 | 5.32E+06 | 3.84E+06 | 3.84E+06 | 0.78 | 0.49 | 4.38 | 4.7 | 5.73 | 6.25 |
| 9243 | 3.26E+08 | 4.18E+08 | 1.02E+07 | 1.09E+07 | 2.30E+06 | 2.69E+06 | 0.17 | 0.13 | 2.06 | 1.1 | 3.91 | 1.12 |
| 9244 | 2.11E+08 | 1.25E+08 | 1.26E+07 | 1.18E+07 | 3.84E+06 | 2.30E+06 | 0.40 | 0.35 | 1.27 | 1.28 | 2.34 | 2.17 |
| 9252 | 6.00E+07 | 1.27E+08 | 2.25E+06 | 1.55E+06 | 2.11E+06 | 2.30E+06 | 1.71 | 1.47 | 2.67 | 3.88 | 7.58 | 3.04 |
| 9255 | 7.44E+07 | 1.04E+08 | 3.97E+06 | 6.19E+06 | 3.07E+06 | 3.07E+06 | 1.89 | 1.40 | 4.79 | 1.94 | 5.86 | 8.14 |

Table S3. **Identical *env* sequences obtained with $Q^2$VOA, NFL and Q4PCR**

| overlap | 9242 | | | 9243 | | | 9244 | | | 9252 | | | 9254 | | | 9255 | | |
|---|---|---|---|---|---|---|---|---|---|---|---|---|---|---|---|---|---|---|
| | $Q^2$VOA | NFL | Q4PCR | $Q^2$VOA | NFL | Q4PCR | $Q^2$VOA | NFL | Q4PCR | $Q^2$VOA | NFL | Q4PCR | $Q^2$VOA | NFL | Q4PCR | $Q^2$VOA | NFL | Q4PCR |
| $Q^2$VOA | 58 | 1 | 4 | 51 | 2 | 1 | 62 | 2 | 1 | 0 | 0 | 0 | 3 | 0 | 0 | 13 | 0 | 3 |
| NFL | 6 | 34 | 1 | 4 | 26 | 0 | 2 | 19 | 3 | 0 | 2 | 0 | 0 | 3 | 1 | 0 | 13 | 0 |
| Q4PCR | 5 | 1 | 37 | 1 | 0 | 8 | 2 | 7 | 8 | 0 | 0 | 2 | 0 | 1 | 3 | 2 | | 18 |
| all shared | 4 | 9 | 4 | 3 | 5 | 3 | 4 | 3 | 2 | 98 | 10 | 21 | 65 | 84 | 95 | 88 | 18 | 22 |

Table S4. **Qualitative analysis**

| ID | sequenced | | MSD | | hypermutation | | indels | | large deletions | | intacts | |
|---|---|---|---|---|---|---|---|---|---|---|---|---|
| | week -2 | week 12 | week -2 | week 12 | week -2 | week 12 | week -2 | week 12 | week -2 | week 12 | week -2 | week 12 |
| 9242 | 204 | 249 | 4 | 4 | 22 | 38 | 23 | 14 | 133 | 169 | 22 | 24 |
| 9243 | 97 | 81 | 2 | 4 | 2 | 2 | 4 | 3 | 80 | 69 | 9 | 3 |
| 9244 | 136 | 110 | 13 | 24 | 2 | 3 | 7 | 9 | 105 | 69 | 9 | 5 |
| 9252 | 133 | 235 | 25 | 131 | 0 | 0 | 56 | 58 | 36 | 39 | 16 | 7 |
| 9254 | 76 | 91 | 0 | 2 | 10 | 5 | 1 | 3 | 22 | 25 | 43 | 56 |
| 9255 | 235 | 185 | 3 | 3 | 14 | 4 | 13 | 8 | 187 | 145 | 18 | 25 |

Table S5. **Probe analysis**

| | probes | Absolute frequencies | | Positive predictive value | Sensitivity |
|---|---|---|---|---|---|
| | | all defects | intact | | |
| all | ≥ 1 | 1595 | 237 | 13% | 100% |
| | ≥ 2 | 1325 | 237 | 15% | 100% |
| | ≥ 3 | 473 | 181 | 28% | 76% |
| | = 4 | 19 | 20 | 51% | 8% |
| ≥ 1 | PS | 518 | 153 | 23% | 65% |
| | gag | 1040 | 197 | 16% | 83% |
| | pol | 1022 | 92 | 8% | 39% |
| | env | 832 | 233 | 22% | 98% |
| ≥ 2 | PS+gag | 258 | 113 | 30% | 48% |
| | PS+pol | 321 | 34 | 10% | 14% |
| | PS+env | 93 | 152 | 62% | 64% |
| | gag+pol | 756 | 78 | 9% | 33% |
| | gag+env | 488 | 193 | 28% | 81% |
| | pol+env | 412 | 89 | 18% | 38% |
| ≥ 3 | PS+gag+pol | 117 | 20 | 15% | 8% |
| | PS+gag+env | 32 | 112 | 78% | 47% |
| | PS+pol+env | 44 | 34 | 44% | 14% |
| | gag+pol+env | 337 | 75 | 18% | 32% |
| = 4 | PS+gag+pol+env | 19 | 20 | 51% | 8% |

Table S6. **Probe analysis for individual participants**

| | probes | 9242 | | | 9243 | | | 9244 | | | 9252 | | | 9254 | | | 9255 | | |
|---|---|---|---|---|---|---|---|---|---|---|---|---|---|---|---|---|---|---|---|
| | | all defects | intact | PPV | all defects | intact | PPV | all defects | intact | PPV | all defects | intact | PPV | all defects | intact | PPV | all defects | intact | PPV |
| ≥ 1 | PS | 294 | 44 | 13% | 117 | 11 | 9% | ND | ND | ND | 48 | 1 | 2% | 59 | 97 | 62% | ND | ND | ND |
| | gag | 75 | 6 | 7% | 138 | 12 | 8% | 97 | 14 | 13% | 325 | 23 | 7% | 67 | 99 | 60% | 338 | 43 | 11% |
| | pol | 287 | 18 | 6% | 8 | 0 | 0% | 2 | 0 | 0% | 340 | 23 | 6% | 43 | 16 | 27% | 342 | 35 | 9% |
| | env | 178 | 46 | 21% | 59 | 11 | 16% | 231 | 14 | 6% | 249 | 21 | 8% | 10 | 99 | 91% | 105 | 42 | 29% |
| ≥ 2 | PS+gag | 36 | 4 | 10% | 116 | 11 | 9% | ND | ND | ND | 48 | 1 | 2% | 58 | 97 | 63% | ND | ND | ND |
| | PS+pol | 230 | 17 | 7% | 8 | 0 | 0% | ND | ND | ND | 43 | 1 | 2% | 40 | 16 | 29% | ND | ND | ND |
| | PS+env | 68 | 44 | 39% | 11 | 10 | 48% | ND | ND | ND | 10 | 1 | 9% | 4 | 97 | 96% | ND | ND | ND |
| | gag+pol | 60 | 4 | 6% | 8 | 0 | 0% | 1 | 0 | 0% | 320 | 23 | 7% | 43 | 16 | 27% | 324 | 35 | 10% |
| | gag+env | 47 | 6 | 11% | 31 | 11 | 26% | 96 | 14 | 13% | 234 | 21 | 8% | 10 | 99 | 91% | 70 | 42 | 38% |
| | pol+env | 84 | 18 | 18% | 2 | 0 | 0% | 1 | 0 | 0% | 249 | 21 | 8% | 2 | 16 | 89% | 74 | 34 | 31% |
| ≥ 3 | PS+gag+pol | 26 | 3 | 10% | 8 | 0 | 0% | ND | ND | ND | 43 | 1 | 2% | 40 | 16 | 29% | ND | ND | ND |
| | PS+gag+env | 8 | 4 | 33% | 10 | 10 | 50% | ND | ND | ND | 10 | 1 | 9% | 4 | 97 | 96% | ND | ND | ND |
| | PS+pol+env | 30 | 17 | 36% | 2 | 0 | 0% | ND | ND | ND | 10 | 1 | 9% | 2 | 16 | 89% | ND | ND | ND |
| | gag+pol+env | 39 | 4 | 9% | 2 | 0 | 0% | ND | ND | ND | 234 | 21 | 8% | 2 | 16 | 89% | 60 | 34 | 36% |
| = 4 | PS+gag+pol+env | 5 | 3 | 38% | 2 | 0 | 0% | ND | ND | ND | 10 | 1 | 9% | 2 | 16 | 89% | ND | ND | ND |

ND, Not Detected
